# Evolution and paradigm shift in forest health research: A review on global trends and knowledge gaps

**DOI:** 10.1101/2024.06.03.597256

**Authors:** Cristina Acosta-Muñoz, Rafael M. Navarro-Cerrillo, Francisco J. Bonet-García, Francisco J. Ruiz-Gómez, Pablo González-Moreno

## Abstract

Forests provide key ecosystem services to human society, and the ability to provide these services depends on their overall health. Forest health is an attractive and interesting concept in forestry research, which environmental, social and political interests have shaped. Assessing forest health is crucial, but finding a single definition of the concept is complex. It is determined by the aim of the forest study, different areas of knowledge, scales of work, technology, methodologies, historical moment or source of funding, among others. With almost a century of scientific evidence, the aim is to identify and contextualise temporal changes in the relevance of this key concept. Trends are analysed through the construction of three main descriptors (state variables, drivers and methods) and the main conceptual subdomains (themes). This review reveals the significant geographical bias in the research, which the Global North predominantly conducts. We observe the evolution of forest health research driven by diverse needs and interests, ranging from air pollution to the multifaceted impacts of climate change. Methodologies applied in this field have also evolved from traditional crown condition inventories to the use of advanced tools such as remote sensing or ecophysiology, improving the characterisation of forest health patterns at both global and individual scales. Forest health research has evolved towards more holistic and multidisciplinary approaches, reflected in the broadening and integration of methodologies and technologies, influenced by historical context, which influence what is being researched today and future scenarios. We identified key knowledge gaps in the scientific literature, in particular the concepts of ecosystem services, Essential Biodiversity Variables (EBVs) and the concept of ‘One Health’. These findings highlight the need for future research to incorporate these critical but often overlooked areas, potentially reshaping future directions and scenarios for forest health research.

## 1. Introduction

The transformation of forests by human activity underscores the imperative need to focus on their preservation and health (1). Forests are essential for sustaining fundamental ecosystem services for biogeochemical cycles and humanity (2). This link between forest health and the capacity of forests to provide such services highlights the importance of understanding and maintaining the ecological integrity of these ecosystems. Recognising and responding on forest health is crucial to ensure their continued contribution to environmental and human well-being. The concept of forest health is an umbrella concept encompassing a wide range of conceptual subdomains (3,4), adopted by practitioners to understand health status (5). This reflects the complexity inherent in the investigation of forest ecosystems, their interaction with human activities and environmental changes. As a result, researchers have used different study perspectives, definitions and research terms over time depending on the focus, scale of work and other aspects considered such as priorities [6], [7].

Researchers have adopted various terms related to forest health such as forest dieback, forest decline or forest decay, associating these processes with the presence of diseases or pests, observing symptoms at tree level (8). At larger scales, forest managers and researchers have traditionally focused on characterizing the potential causes and spatio-temporal patterns (9). Thus, the terms reflect not only tree mortality, but also a general loss of vigour and yield that is spread over relatively large areas and is often related to high environmental stress (10). Recently, the term forest health has evolved including also structural and functional aspects (11). For instance, some authors define a healthy forest as one that includes a mosaic of successional patches representing all development stages (12). At the same time, other more holistic terms such as forest condition, forest state and forest integrity have been proposed. Among them, forest integrity has been one of the latest suggestions, defining the overall capacity of a forest system to sustain composition, structure and function within the historical range of variation (13,14).

Beyond the different conceptual subdomains we have listed above (used to define or assess forest properties), we argue that the science and research has changed over time, with different themes and associated terminology. We determine significant changes in recent decades based on the following three main descriptors: a) attributes to measure forest condition; b) drivers impacting on forest health condition (i.e., biotic or abiotic); and c) technologies and methodologies associated with measurement and analysis of the two previous aspects.

With the expansion of science, the ever-deepening knowledge and the rapid pace of publication seen in almost all scientific disciplines (15), this review emphasises the critical importance of understanding what has brought us to the present. Understanding the historical context of the discipline lends depth to current perceptions of forest health and is crucial to addressing the challenges ahead. We not only value the foundations of our knowledge, but also recognise the importance of following a deliberate and informed path for the future and innovation in forest ecosystem research.

The inherent dynamic of constantly evolving research approaches is also related to changes in the methodologies and technologies. The most frequent measurements have been crown condition and tree damage (e.g. defoliation and discoloration), or growth in terms of biomass and diameter increments (16,17). More holistic approaches go beyond the tree level to characterize population, community, and ecosystem properties such as biodiversity and regeneration dynamics. Regarding the drivers, forest health is affected by several disturbance agents of different origin, and which can impact forest systems in a complex and interactive way (11,18). In addition, context-dependency of the relevance of different abiotic and biotic agents affects the overall research outputs, with bias towards scientist’ geographical regions and specific taxa (19).

Forest researchers have put enormous effort into forest observation and monitoring to understand forest health in relation to forest condition and related drivers (20). These encourage the development of a wide range of methodologies aiming to characterize forest ecosystem trends to inform policy and management decisions. These methodologies include direct measures of vegetation such as physiological (e.g. photosynthesis, pigments, water transport, respiration), structural measures derived from traditional forest inventories (e.g. growth, dendrochronology), measures related to external agents, but with implications on vegetation (e.g. drought, changes in land cover and land use), or measurements related to the role and functioning of forest as an ecosystem (e.g. nutrient cycling and productivity) (9,21). Lately, an increasing relevance of measurements derived from remote sensors deployed at satellite and unmanned aerial vehicles has been observed for the detection of non-visible phenomena in the forest (22–24). The convergence of these advanced methodologies together with technological innovation allows for an ever deeper understanding of forests and their dynamics.

Searches in the main scientific information databases (e.g. Web of Science or Scopus) show that current knowledge on forest health is fragmented across several research disciplines (forestry, environmental sciences, ecology, entomology, plant sciences, remote sensing, biodiversity conservation, geosciences, agricultural and biological sciences, earth and planetary sciences, social sciences, computer sciences, biochemistry, genetics and molecular biology, and others). Several attempts have been made from different disciplines to review and describe these changes in forest research (25) and to synthesize the conceptual frameworks around forest health (11) without having a complete picture of the temporal dynamics of the concept. Systematic and bibliometric reviews of scientific literature is key to synthesize a research field and to understanding the conceptual trend. We used this approach to understand the temporal and regional trends, research conceptual subdomains and methods used on forest health assessment and monitoring at global scale.

Exploring the evolution of approaches to scientific research in forest health, to better understand trends, developments and challenges, and the implications this has for the way science is conducted globally, is of relevance [8]. In our study, we aimed to i) contextualise the status of and approaches used in forest health research (i.e. scientific output, main contributors, issues and keywords), ii) assess the temporal evolution of recurrent terms in forest health research (i.e. a complete temporal map of relevant keywords), and iii) understand temporal trends in the main conceptual subdomains encompassing forest health (i.e. topics) and the three descriptors of forest health introduced in this study: condition (variables used to measure the state or condition of forests), drivers (abiotic or biotic agents causing changes in forest condition) and methodologies (techniques used to assess forest health).

We conducted a scientific literature search in academic databases covering a wide spectrum of forest health terminology (26). Data mining was applied to extract information and patterns from large bibliographic datasets that qualitatively, quantitatively and graphically allow a deeper understanding of scientific production (27–29). Using a systematic review, we contributed to temporally characterise the forest health concept to provide a holistic definition. Finally, we discuss the gaps and future potential conceptual subdomains and descriptors that seem to arise in the research field.

## 2. Material and methods

The workflow carried out for the analysis included (Fig 1): data collection, scientometric and bibliometric analysis and visualization (all of these are detailed below). Specifically, to generate the results of the first objective we performed a descriptive analysis of scientific production, a bibliometric analysis of maps in terms of co-occurrence of keywords, and an analysis of publications and contributions. To achieve the second objective, we provided a time trend analysis of recurrent terms. And for the third objective we performed a temporal trend analysis on a semantic clustering of the keywords (obtained from the previous objectives) in the three forest health domains established in this study: forest condition, drivers and methods.

**Fig 1.**
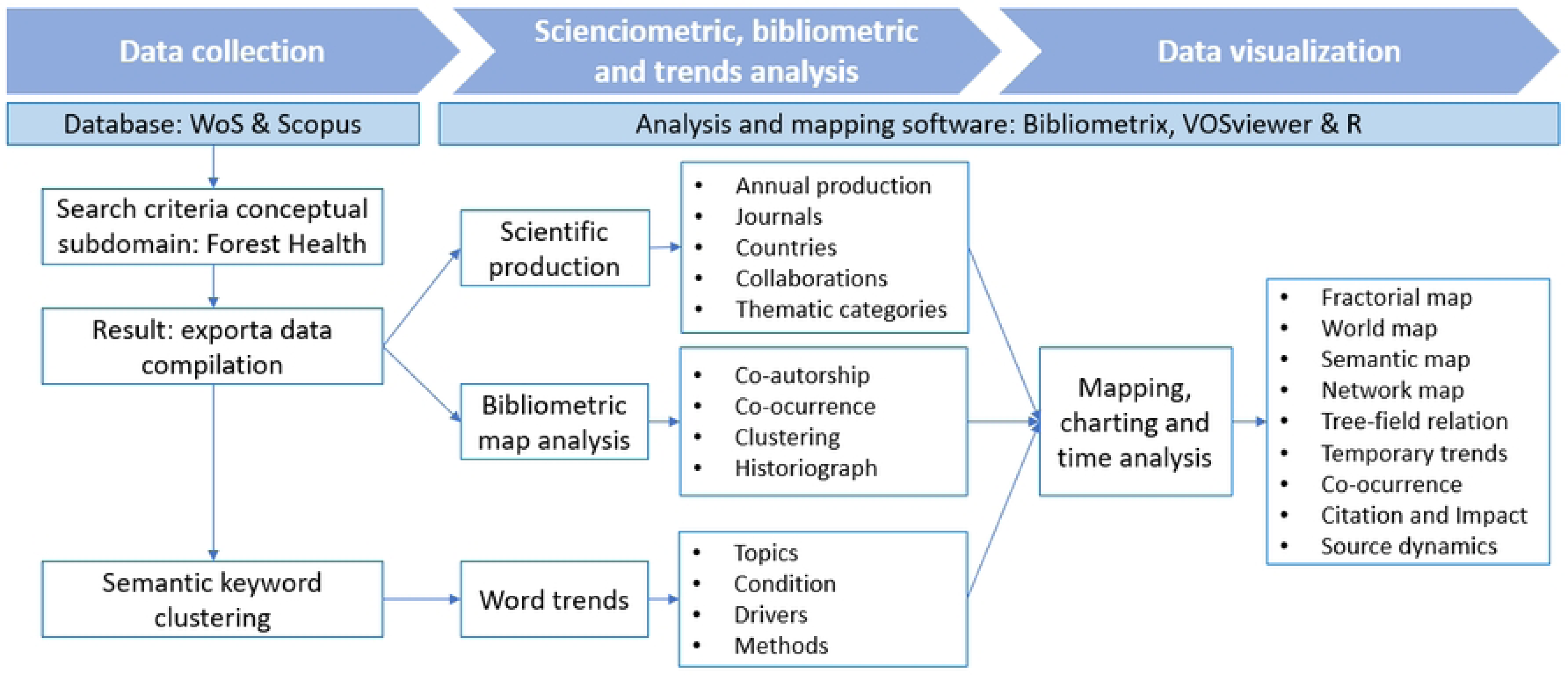
Graphical summary of the bibliographic and bibliometric review workflow for the study of forest health concepts evolution.

### 2.1. Scientific Data Base Search

The Web of Science (WoS) and Scopus database was chosen as high-impact search engines for formal scientific publications, excluding non-conventional literature (grey literature). In January 2024, we carried out a preliminary search with a single keyword linear strategy using “forest health” in Title, Abstract and Keywords fields, for all recording times in the database. The temporality of the search was not limited, as the aim was to know completely all the existing records in the databases from their origin to the present. The search was refined to include only disciplines related to biological, environmental, forestry, earth sciences or methodological sciences, excluding humanities or medical sciences.

Titles and abstracts of the 100 most relevant articles from each year were read and reviewed for the state of the art of forest health research and scientific production. Potential articles that could contain definitions or different terminology of forest health were read in detail (i.e. reviews and highly cited papers). From this preliminary review, a list of conceptual subdomains for the umbrella term forest health was compiled and used in the final database query.

The following search strategy was used to obtain the corpus: Title / Abstract / Keyword = [“forest health” OR “forest mortality” OR “tree mortality” OR “forest integrity” OR “forest state” OR “forest decline” OR “forest decay” OR “forest dieback”]

### 2.2. Descriptive analysis of the status of forest health research

The records obtained from the final query were analysed using the quantitative bibliometric analysis algorithms of the R package *Bibliometrix* and the associated application *Biblioshiny* (26). First, we conduct a descriptive analysis that characterises the scientific production over time, main contributing authors, the co-authorship network, country publication impact, geographically contextualising the main journals and funding agencies. Secondly, Sankey diagrams were used to focus the analysis on keywords to identify patterns and trends in the main terms used to describe forest health research in the above context (30).

### 2.3. Temporal evolution in forest health issues

An analysis of relevance and development in research topics based on the initial systematic review separated the selected keywords into four main groups: a) “motor themes”, that is, themes well developed and important for the structure of the research field, b) “emerging or declining topics” when they are both weakly developed and marginal, c) “basic and transversal topics” which are important for a research field but are not developed and d) “niche with a specialized character”, which are peripheral and specific topics for the research field.

Temporal changes in the concept and application of forest health were assessed by an automatic keyword network analysis using WOSviewer (29), based on the previous review and debugging of retrieved words. We used co-occurrences of keywords that appeared together in the title, abstract or keyword list, and that were mentioned at least 10 times, giving a total of 1,731 keywords. We then plotted the top 1,000 keywords in a network and recurrence map. Finally, this network was overlaid with the year of publication to identify temporal trends in keyword association.

### 2.4. Clustering of conceptual sub-domains and temporal trends

We also implemented an approach considering a manual semantic keyword clustering to understand temporal trends across the three domains of forest health considered in this study (e.g., condition, drivers and methods) and main topics or definitions of forest health. From the bibliographic search and subsequent download of references, for each year, the 50 most relevant author and recommended keywords were extracted (keyword PLUS - index of terms automatically generated from the titles of cited articles). This set of keywords went through a process of cleaning up duplicates, normalising or removing special characters, reviewing compound words and reducing some words to their basic roots. From this list of keywords for all years, duplicated words were removed and grouped semantically. These terms were classified into descriptors of topic (theme or discipline in forest health concept), condition (i.e., variables used to measure forest state or condition), drivers (abiotic or biotic agents causing changes in the forest condition) and methodologies (techniques used to assess forest health). Within each descriptor type, we grouped the terms according to similar semantic meaning (Table S1 Supplementary Material). In addition, when reviewing the list, other words were proposed for being particularly relevant for the analysis based on the preliminary review indicated in section 2.1. For each term, we compiled their occurrence in the abstracts of each of the scientific records retrieved and calculated their frequency per year. All analysis were carried out with R version 4.3.2. (31).

## 3. Results

### 3.1. Descriptive analysis of scientific production in forest health research

#### 3.1.1 General findings

Scientific evidence in forest health from 1934 to 12/2023 shows an exponential publication trend (as in most scientific disciplines), with an annual growth of 7.8%, although there was a decline during the COVID-19 pandemic (Fig S1; Table S2 Supplementary Material). We analysed 10,338 papers from 1,511 sources, with 26,025 authors, highlighting that the 20 most prolific authors account for 9.37% of the publications, indicating that forest health research is highly diversified in terms of researchers involved in this topic. The top 4 authors stand out, with more than 60 publications each (Fig S2 Supplementary Material). The leader in publications, Camarero J.J., uses growth ring analysis and remote sensing to study the interaction between forests and their environment, underlining the importance of longitudinal studies for conservation policies. The first recorded publication was by Veblen T.T. in 1983, focusing on forest instability and tree mortality using dendrochronology. The most cited authors are Allen D.C. and Breshears D.D. in 2010, for investigating global forest decline and drought.

#### 3.1.2. Publication impact countries

Considering the 10 countries with the highest publication record on forest health, USA marks a significant difference with the rest of the world throughout the whole period studied (Fig S3 Supplementary Materials). Next, Germany was the country with the longest track record in related research. Canada increased its relevance in last decades, overtaking Germany in 2007, and reaching the second position. Since 2010, China obtained an exponential increase in the number of publications, reaching currently the third top position.

In terms of cross-country collaborations, 5 clusters were observed. A first cluster (Fig S5 Supplementary Materials) related by spatial continuity and ecological similarities, including countries in North America (USA, Canada, or even Mexico); they in turn also related to other countries by their latitude (Russia), or by the problems raised and methodological challenges to cover large countries (such as China or Australia). A clear cluster was observed where South American countries seem to be very closely aligned and related to the United Kingdom, the Netherlands and Japan. The last distinct cluster related Eastern and Northern European countries to New Zealand.

Despite the bibliometric reflection, the literature review shows that the geographic origin of affiliation of the main authors of the publications does not determine the area of study in the research. Collaborative research and international co-authorship favours diversification of the study regions addressed beyond their own borders. Origin of funding agencies was largely consistent with the most productive countries (Fig S6 Supplementary Materials).

#### 3.1.3. Research approaches and issues

The search returned 13,677 keywords proposed by authors and 10,399 KeyWords Plus. The Sankey diagram (Fig 2) shows boxes of different sizes and colour intensities allowing to identify the areas of greatest activity and connection, among the 10 most common keywords (themes), the 10 countries with the highest scientific output and the top 10 thematic journals (Fig S7 Supplementary Materials).

**Fig 2.**
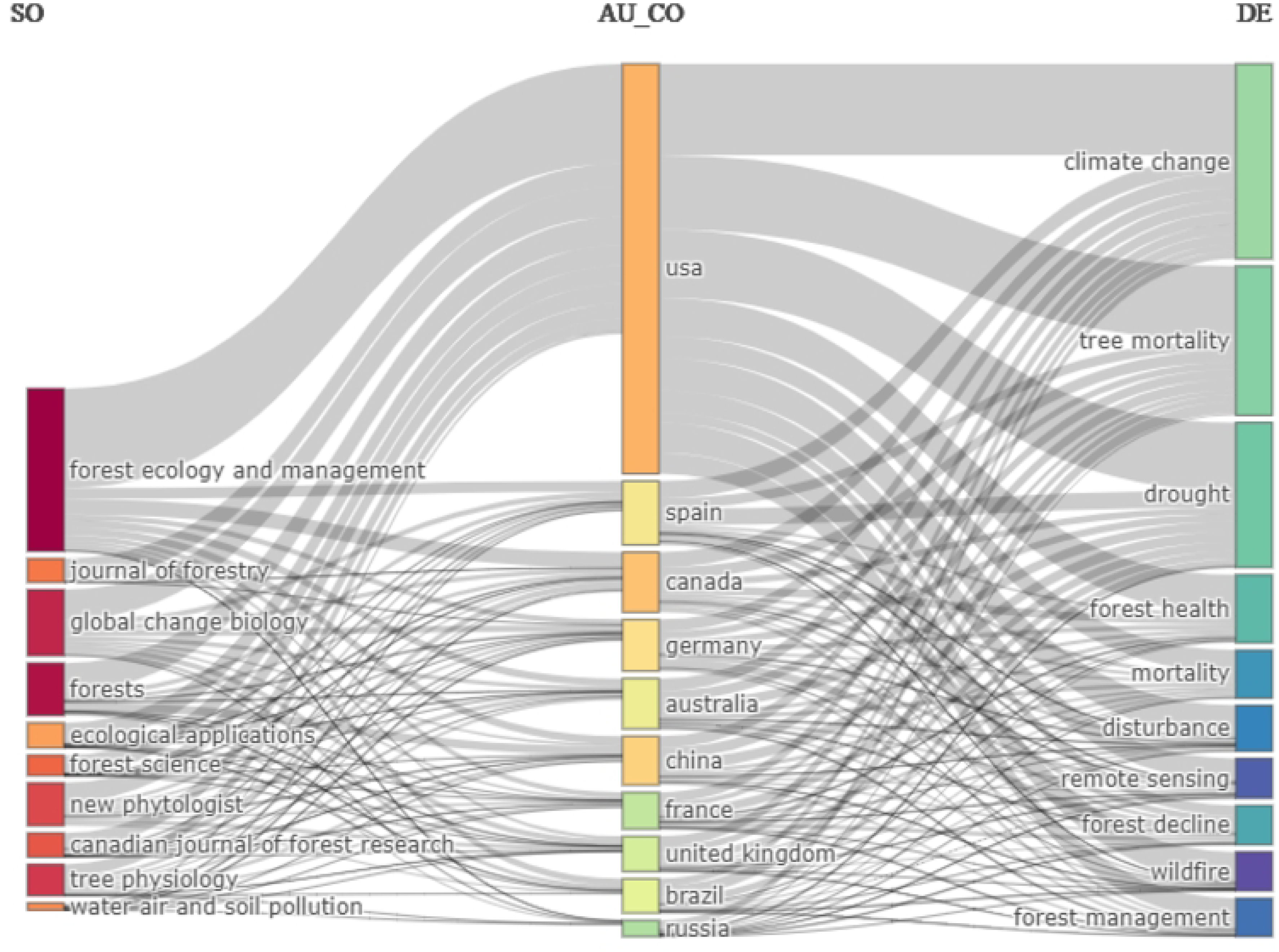
Sankey diagram showing the relationships between frequent journals (left) from the top 10 publishing countries (middle) and the top 10 most mentioned keywords (right) in forest health-related research.

The results highlight that the top countries publish most frequently in journals related to forest management and also ecology. There are variations in keyword priority between countries, with a general focus on climate change, although Spain and Australia also highlight drought.

Co-occurrence analysis showed the 50 most frequently mentioned keywords (Fig S9, supplementary material), revealing “mortality” as the most common term, linked to climate change and drought, and associated with forest vulnerability, climate responses, forest health and growth. Clusters were identified focusing on forest decline due to stress, nitrogen deposition and soil problems, specifically related to pine and spruce. Another cluster emphasises forest management dynamics, impacts and biodiversity. A final cluster addresses the effects of fire and pests on specific species and sites.

### 3.2. Temporal keyword analysis

We grouped the keywords on the x-axis showing the relevance of the topics and on the y-axis the degree of research development (Fig 3). We found that the core group of most relevant topics contains research related to ‘climate change’, ‘tree mortality’, ‘drought’ and ‘fire’. Another group of core themes, although less detailed (compared to the previous one), are ‘forest health’, ‘remote sensing’, ‘dendrochronology’, ‘bark beetle’ and ‘biodiversity’. Of the relevant core themes with the highest level of development, the keyword ‘disturbance’ is the most developed, followed by ‘forest management’, ‘wildfire’, ‘prescribed fire’ and ‘Pinus ponderosa’. The results show that in general, the most developed topics with a high level of specialisation are those related to the physiological processes of the forest, containing keywords such as: ‘water stress’, ‘hydraulic failure’, ‘photosynthesis’, ‘carbon starvation’. Finally, in the group of underdeveloped or unused keywords was ‘forest decline’ when talking about more concrete process words, as well as ‘atmospheric pollution’ and ‘ozone’, issues that were relevant but already little studied.

**Fig 3.**
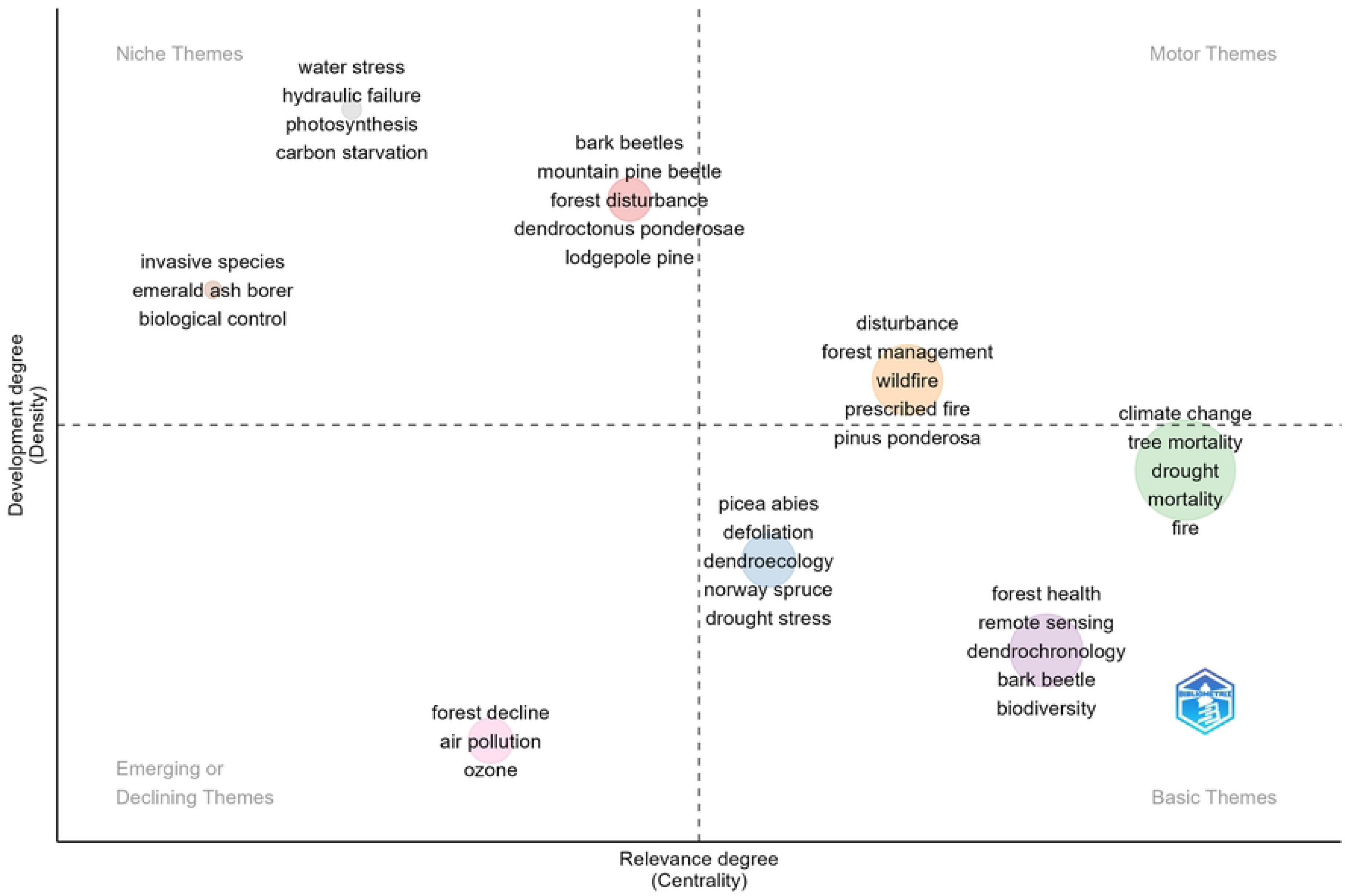
Word grouping map by themes of relevance and development in research on forest health: a) “motor themes”, well developed and important for the structure of the research field, b) “emerging or declining topics” weakly developed and marginal, c) “basic and transversal topics” important but not developed and d) “niche with a specialized character”, peripheral and specific topics for the research field.

The temporal network analysis of the top 1000 keywords showed a total of four clusters over the study period (1934-12/2023) (Fig 4). Although the entire period analysed was included in the graphical representation, due to the low number of publications recorded before the mid-1980s in the databases consulted, the first cluster (purple colour) appears from this initial stage of scientific production. It was closely linked to the concept of “forest decline”, especially in topics related to “air pollution” (“nitrogen”, “ozone” or “acidification”). Around late 1990s and early 2000s, the diagram showed a second broad cluster (in blue-green) highly associated with the term “growth” and “forest health”. These terms were mostly related to monitoring and development-related terms (e.g. “stands”, “competence”, “deforestation”, or issues related to “biomass” and “carbon sequestration”). Approximately in 2010, a new cluster (in green) appears with quite wide range of terms of similar importance but related to ecosystem processes and characteristics: (e.g. “biodiversity”, “dynamics”, “disturbance”, “management, “conservation”, “restoration” and topics associated with “fire ecology”). This cluster seem to converge into the concept of “tree mortality”, peaking around 2015. Finally, in the most recent period (in yellow), the research activity focused on “climate-change”, showing a great interest in the variables measured and the tools related to “ecophysiology” and “remote sensing”.

**Fig 4.**
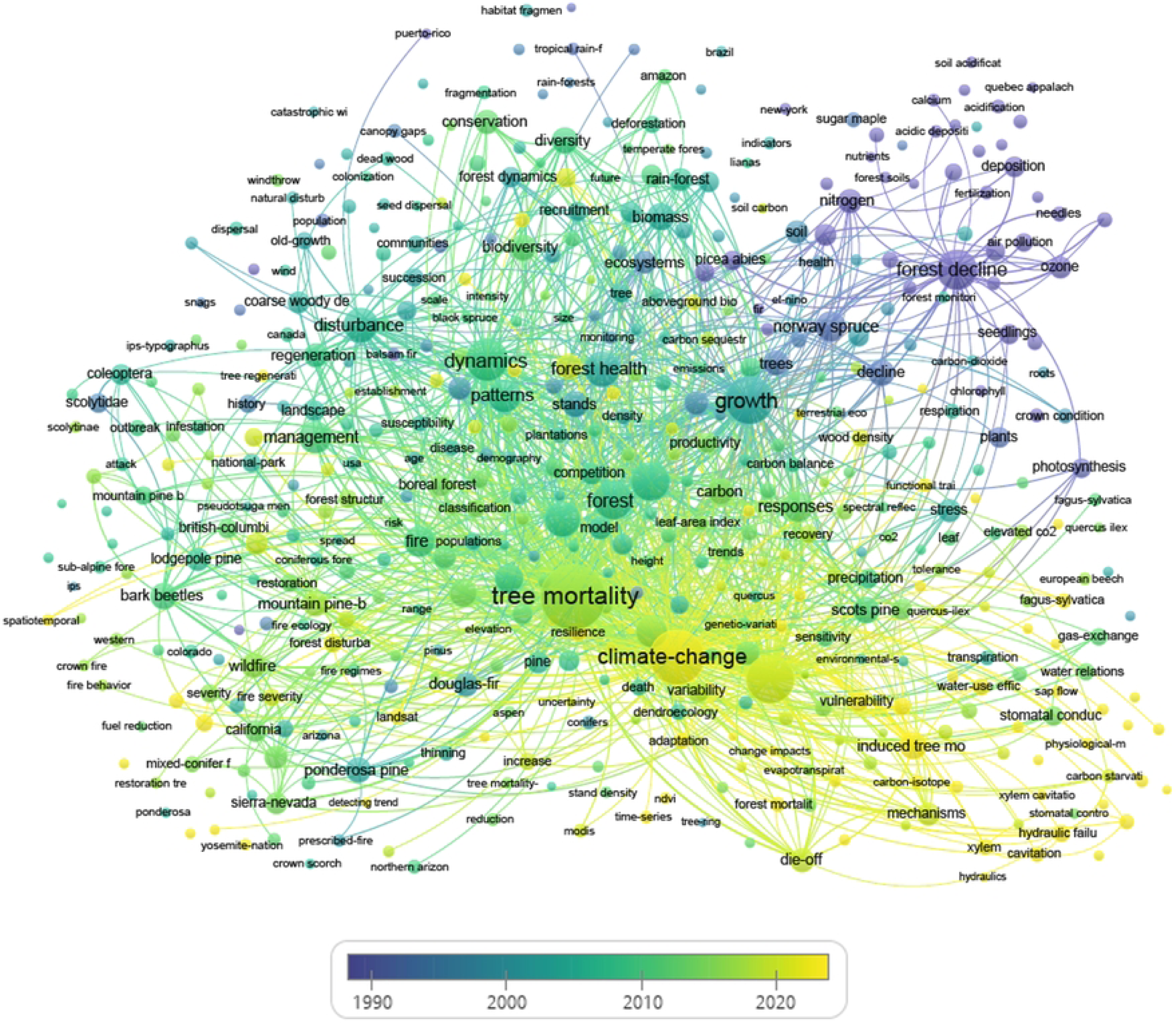
Temporally normalised co-occurrence of most frequent keywords related to forest health research (VOSviewer graph).

### 3.3. Clustering and trends in forest health descriptors

Manual classification of keywords in forest health reveals four sets of issues clustered into the four established categories: topic, condition, drivers and methodologies (Fig 5), with fluctuations in the 1970s to mid-1980s due to low production and thematic dispersion (removed from the graphical representation), stabilising since the 1990s. Interest in “forest decline” has declined, being replaced by “tree mortality”, and “forest health” had a peak around 2005, followed by a recent decline in the last decade. The attributes measured to characterise forest condition reflect evolving environmental concerns, from “pests and diseases” and “air pollution” to a growing interest in “climate” and “fire”, which have become a constant concern. Methodologically, understanding the functionality of organisms based on “ecophysiology” has declined in relevance, while tools such as “inventories” remain constant. Since the 1990s, the use of “Geographic Information Systems” and “remote sensing” has grown significantly, just as modelling-based methodologies have increased in importance in recent years, although to a lesser extent than the former.

**Fig 5.**
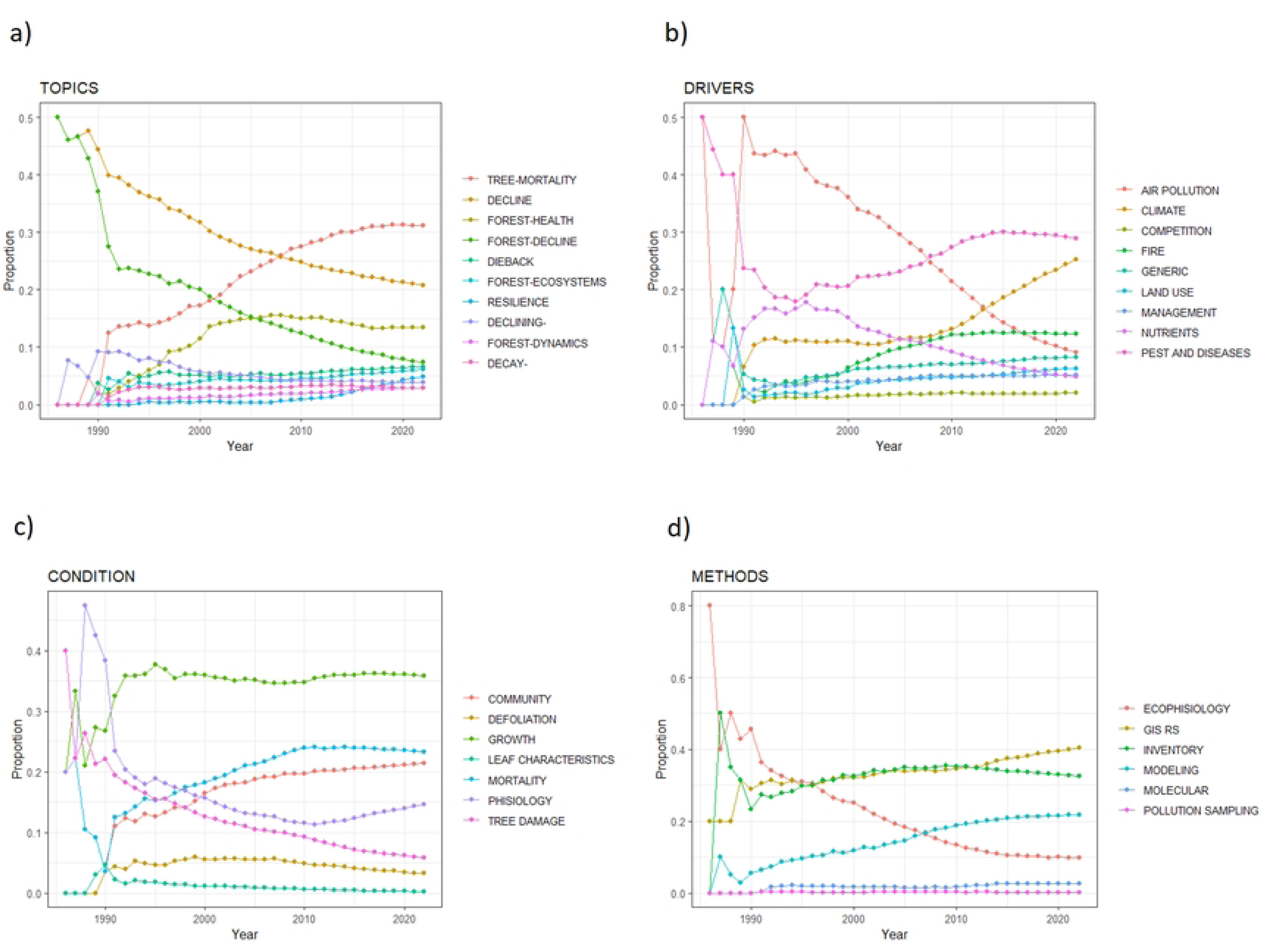
Proportion of occurrence of keywords per year considering four different aspects of forest health: 5a) topic (theme or discipline in forest health concept), 5b) condition (variables used to measure forest state or condition), 5c) drivers (abiotic or biotic agents causing changes in the forest condition) and 5d) methodologies (techniques used to assess forest health).

## 4. Discussion

Over the last 90 years, we have seen a remarkable increase in the publication of research on forest health, which underlines the need for an assessment of its evolution. The study synthetically contextualised the accumulated body of knowledge, identifying substantial changes in the ways in which forest research is approached, studied, measured and the technologies associated with it. The results obtained showed aspects related to how science has been done so far, detecting new emerging concepts or trends. Particularly noticeable are the changes in scientific production, together with variations in the concepts and methods employed by the forest health research community, reflecting how these aspects have been linked to the main environmental concerns of each period, to funding or to the geographical area of influence of the researchers.

Following this bibliographic and bibliometric review, we aim to further explore the rationale for temporal trends, highlighting their importance and implications at each point in time. The study allows us to identify both significant advances and gaps in knowledge, contributing to the configuration of new lines of research that respond to emerging challenges in forest health.

### 4.1 Scientific production on forest health

As with many other concepts in various disciplines (32,33), research on “forest health” is experiencing exponential growth in terms of the number of articles published. Leaving aside the general trend in academic production, the exponential growth observed denotes that the concept under study remains dynamic and continues to be of interest to researchers. This vigorous increase in the production of scientific literature reflects not only the growing global concern for the state of our forests but also the recognition of the complexity and multidimensionality that characterises them (34). As such, there remains a need for a collective effort to understand and mitigate the impacts of threats such as climate change, tree diseases and deforestation (35,36).

The affiliation origin of the researchers revealed important information on how the knowledge is created regarding the topics about forest health. Our findings show that USA, Canada, and Germany are the countries of affiliation origin for most of the researchers working in the target topic. These results unveil two important biases:

First, there is a mismatch between the most publishing countries and those which harbour a higher percentage of forests worldwide. According to the latest Food and Agriculture Organisation (FAO) report on global forest resources (37), more than half of the world’s forests are concentrated in 5 countries: Russia (815M ha - 20%), Brazil (497M ha - 12%), Canada (347M ha - 9%), USA (310M ha - 8%) and China (220M ha - 5%). Only Canada and the USA are among the countries with large areas of forest with substantially more forest health articles compared to the others.

Second, there are not countries located in the tropics in the list of the higher publishing records. This situation is more remarkable if we consider that the largest percentage of forest (45%) is in tropical areas. This uneven distribution of publications between the Global North and the Global South has been described in other disciplines such as ecology (38): most of the research is done in the Global North although both the biodiversity and the forests are mainly in the Global South. This geopolitical situation impacts very deeply in the completeness of the “forest health” concept since it does not consider the views of researchers from the countries with more forest cover.

### 4.2 Evolution of the most relevant keywords on forest health research

#### 4.2.1. Integrating Keywords: Uncovering patterns

The total number of keywords found in “Climate change”, “tree mortality” and “drought” were the topmost common keywords mentioned in forest health related research (Fig 3 and S9 Supplementary Materials). In fact, a large set of forest health studies have built on the delicate situation of forest ecosystems worldwide with large-scale mortality processes driven by climate drivers (9,39). Interestingly, the relevance and development analysis considered these terms as “Basic Theme” showing a high relevance and a medium degree of development, which indicates their current popularity but also further room for development compared to themes such as disturbance or wildfire.

In this “Basic Theme” group, the analysis also highlighted terms such as “remote sensing”, “defoliation”, “dendrochronology” and “biodiversity”, revealing a multidisciplinary and multi-scale approach to capture the complexity and dynamism of forest ecosystems. This approach demonstrates a broad perspective on integrated, multi-scale forms of forest measurement: such as defoliation as a measure of forest response at the leaf level, dendrochronology as a measure of growth rate at the tree level, the use of remote sensing allowing extensive monitoring of forests at the landscape level, or biodiversity as a manifestation of forest structure and functioning at the community level (40–42).

Among the “Niche Themes” (high density and low relevance), we identified three main groups that seem rather peripherical or with regional interest to the research field. From the initial reading and bibliographic review, we found that the research field of invasive species and beetle outbreaks mostly concentrated in North America on conifer forests (43–45) and on the other side pure ecophysiological studies (46). The former group reinforces the idea of the bias toward the Global North: *P. ponderosa* is a heavily timbered species typically found in temperate areas of North America (47). Besides, concerns about bark beetles and prescribed fire are a management activity also frequently used in temperate areas of the Northern Hemisphere (48). Regarding the “Emerging or Declining Themes” with low development and relevance, it is remarkable how the clustering process identifies forest decline, pollution and ozone as themes that are no longer mainstream regarding forest health. These topics refer mainly to the events of acid rain that were relatively common in Europe and North America during the second half of 20th Century and even nowadays in China (49).

### 4.2.2. Origins and context of paradigm shifts

The temporal change in the proportion of keywords tells a history very useful to understand the research topic of “forest health”. The analysis of the temporal evolution of keyword clusters reveals two main patterns (Fig. 4): a) there is a consistent trend towards a higher level of knowledge integration across the time series and b) there is a clear link between the evolution of global research and environmental challenges at each point in time and the changes in forest health research. Based on this analysis, we identify four different temporal clusters that have occurred sequentially (Fig.6):

*I) The arise of global environmental problems linked to atmospheric pollution.* At the beginning of the time series, we found monocausal approaches to forest health disturbances, where the most important drivers were “pests and diseases” as well as “air pollution”. This earliest cluster contains concepts, which are attributable to well defined scientific disciplines: “nutrients”, “forest soils”, “fertilization”, “ozone”, “seedlings”, “calcium”, etc. This pattern indicates the low level of discipline integration that the target concept experienced prior to 1990. Furthermore, it shows clear links with the first modern environmental movements worldwide took place between the 1960s and 1970s, focusing on nature conservation and environmental protection. Predecessor events are the book “Silent Spring” by Rachel Carson (1962), which denounced the harmful effects on the environment of the massive use of chemicals such as pesticides. The first “Earth Day” (1970), the United Nations Conference on the Human Environment in Stockholm (1972), and the “Energy Crisis of 1973” awakened awareness of the dependence on oil and the search for alternative sources. A central forest health topic in this cluster is “forest decline” with strong links to acid deposition, air pollutants, and ozone. In fact, “forest decline” was a terminology commonly used to depict the research concern about forest deterioration due to air pollution mostly in Northern Europe and North America (51). This forest problem gained international relevance in the 1980s with “The Geneva Convention on Long-Range Transboundary Air Pollution” (1979), “The Vienna Convention for the Protection of the Ozone Layer” (1985) or the signing of “The Montreal Protocol on Substances that Deplete the Ozone Layer” (1987). This environmental problem kept research strongly active until the end of the 20th century. It is from the 1990s onwards that the evidence on the effects of pollution began to be related to human health and ecosystems, and although the global burden of pollutants has been increasing in the first two decades of the 21st century, efforts are being made to continue reducing them (49).
*II) Global environmental conservation.* The second cluster is dominated by the decline and physiology of the forest, appearing in late 1990s and early 2000s. Thus, the methodologies mainly used are related to ecophysiology and forest inventories. This period shows concepts with a higher level of integration among disciplines and knowledge bodies: “growth”, “competition”, “forest health”, “ecosystems”, “carbon sequestration”, etc. It also reflects the arise of current environmental problems such as carbon emissions and deforestation. This may be mainly due to the events that took place during the 1990s, where concerns with a more holistic and multidisciplinary view of nature conservation and the environment began to broaden. At this time, among others, the most famous world summits took place: “The United Nations Conference on Environment and Development” or better known as “The Rio de Janeiro Summit” (1992), laid the first foundations for the signing of the United Nations Framework Convention on Climate Change and the signing of the treaty “The Convention on Biological Diversity”, being the first global agreement to promote aspects of international cooperation in the conservation and sustainable use of biodiversity. The famous “Kyoto Protocol” (1997) on the reduction of greenhouse gases that cause climate change is also approved in this period. Almost simultaneously, the FAO publishes a report that highlights the deforestation of large tracts of tropical forests in Latin America, Africa and Asia (52). At the same time, the achievements of the international policy on air pollutants reduced to some extent the pressure of air pollution on forest ecosystems (53) and consequently in the forest health research field. In summary, this temporal cluster represents the initial foundations for a more holistic and larger-scale view of the planet’s global problems, evidenced among the key words in scientific publications of the time, leading to the use of more multidisciplinary, integrative and comprehensive concepts.
*III) Multi-*causality *and tree mortality.* The research that begins with the 21^st^ century shows a multi-causal thinking in the problems that occur in the deterioration of forests and the environment. This group shows a wide range of concepts, where the words that stand out the most are “patterns”, “dynamics” and “disturbances”. Now the forest problems are based on multi-causality, a more complex vision that can be studied not only at the tree level but at different scales and in a multidisciplinary way. The characterization of ecosystem dynamics is based on classification, offering scales, intensities or patterns that measure diversity, fragmentation, deforestation, succession, competition, susceptibility, regeneration, among other processes (54–56). Furthermore, other concepts such as “management” and “restoration” also emerge as a key concept suggesting a more applied vision in the forest health research agenda (57). At the end of this period, the research agenda converge on the topic of “tree mortality” with numerous links to a wide range of concepts. Other highly integrative concepts also appear (e.g. restoration, resilience) reinforcing the paradigm of multi-causality in forest health research (58,59). “Wildfires”, its consequences, and some methods used to monitor them, are present here to explicit the environmental issues addressed in that time. It is no longer enough to quantify the causes of disturbances in the system, but rather the effects of the disturbances themselves, in search of solutions and to assess both the damage and the improvement in the global balance.
*IV) Climate change driven-research.* The most recent cluster contains mainly concepts related to “climate change” (e.g. “vulnerability”, “adaptation”, “change impacts”, etc.). Interestingly, this cluster seems to reduce the degree of knowledge integration as scientists are focusing mostly on understanding the consequences of climate change on forests; although this challenge is much more complex than those described previously. This is evidence of the greater environmental awareness, both social and political, in the mitigation of climate change. One example is the approval of the “The Paris Agreement” (2016), which establishes a global framework on climate change focused on concrete aspects such as curbing global warming and achieving carbon neutrality before the end of the century, where the use of the best available science and technology are directly included to improve the conditions of the planet. In fact, “climate change” can be considered a so called wicked problem (60): multifaceted problems with fuzzy definition, elusive and complex solutions. This explains why the current distribution of words within the drivers becomes more equative. It is also the boom in technological development that derives part of these efforts in generating instruments, methods and measurement and evaluation techniques that are increasingly more accurate, reliable, and accessible. The emergence of portable electronic equipment or geospatial technologies like remote sensing, were a breakthrough to obtain continuously and efficiently data across different spatio-temporal scales. This idea is supported by the presence within this cluster of methods of assessment (e.g. carbon-isotope, dendroecology) and a great amount of forest condition variables (e.g. evapotranspiration, water-use efficiency, stomatal conductivity, hydraulic failure, etc.) currently measured with sophisticated ecophysiological sensors (e.g. “gas-exchange” related to Eddy covariance towers or photosynthesis sensors).

**Fig 6.**
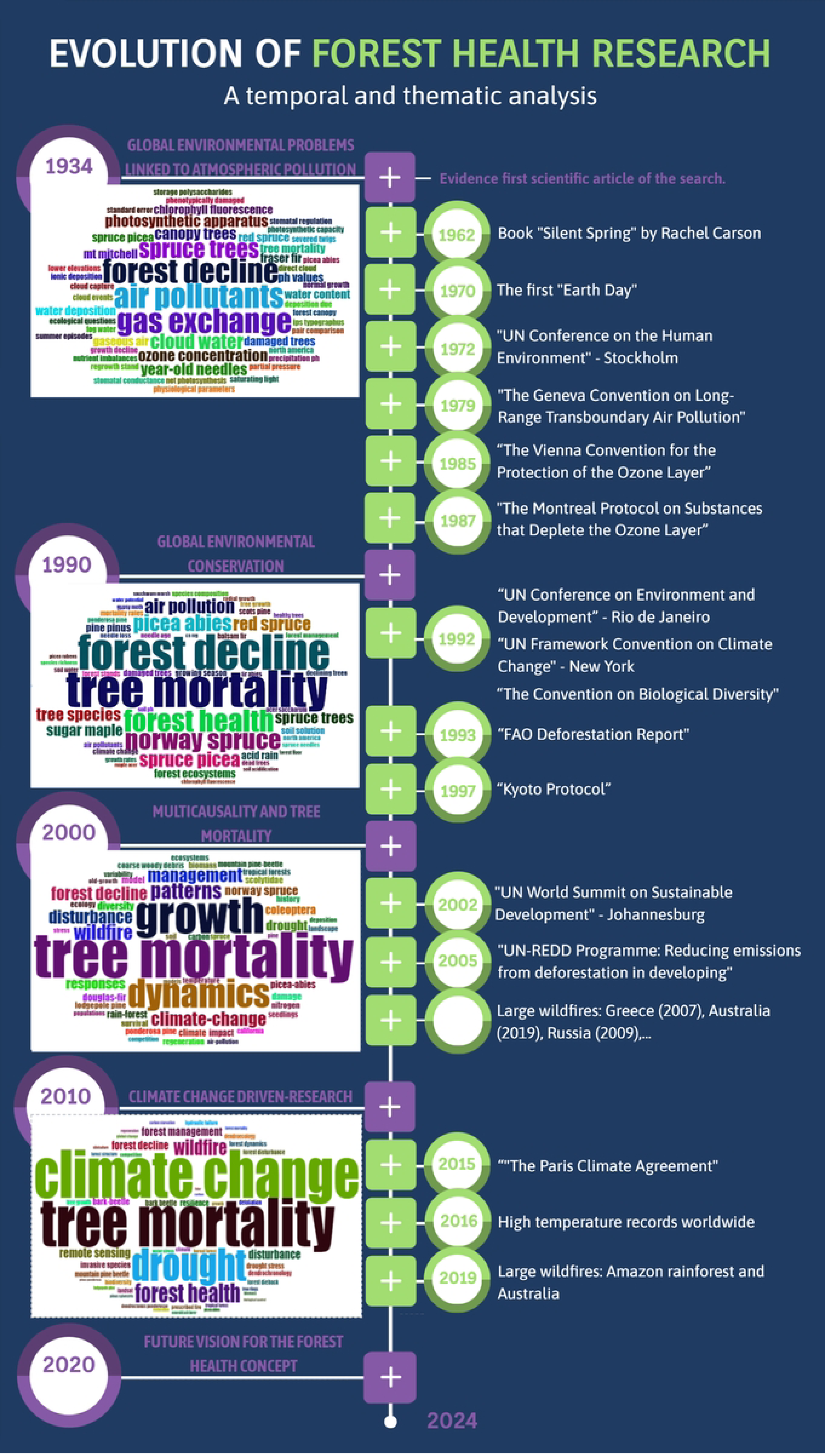
Historical and thematic analysis of nearly a century of advances in forest health research, illustrating key international environmental and socio-political milestones (to the right of the figure) that align with shifts in prevailing themes and scientific terminology (evidenced by the word clouds to the left of the figure).

### 4.3 Trends topics, concepts and methodologies on forest health research

We also found similar temporal patterns from the temporal analysis of the four different descriptors (topic, condition, drivers, and methodologies; Fig 5). First, we have identified a temporal trend towards higher complexity. In the case of drivers, there is a clear trend from monocausal to multicausal drivers of interest. From the predominance of “air pollution” and “pest and diseases” during the first decades, to the emergence of other concepts: “competition”, “land use”, “management”, and “fire”, which ultimately end in steep increase of climate related drivers related to the “climate change” paradigm (Fig 5C). This could mean that the drivers of change in the “forest health” research domain are more complex now than they used to be decades ago. A similar pattern can be found in the group of “topics” words (Fig 5A). The predominance of forest decline leads to a richer scenario where tree mortality, forest health and still forest decline have certain importance. Regarding the condition words, the last decades show also the rise of terms that imply a richer and more integrative approach: community, mortality, growth, etc (Fig 5B). They coexist in a scenario more equitable than the existing at the beginning of the time series.

On a second note, methods have followed the divergence and increased in complexity of the other forest health aspects (Fig 5D). Finding appropriate methods to measure forest condition has been always a major challenge in forest health research. Different types of methodologies and techniques to assess forest status have been continuously evolving. Historically and up to the present day, classical inventories have been an objective measure of forest species composition, quantity and distribution of trees, as well as tree quality based on simple structural measures (61,62). These inventories have become more complex as measurement tools have evolved, although in general they are techniques that require little instrumentation, they are limited in the amount of land that a team of people can cover. In this sense, important forest monitoring programmes emerged in the 1980s.

Some of these programmes are the International Co-operative Programme on Assessment and Monitoring of Air Pollution Effects on Forests (ICP Forests) of the United Nations Economic Commission for Europe (UNECE), since 1985 (63); or the Forest Health Monitoring (FHM) of the United States, which began in 1990 (64). Many of these inventories were accompanied by physiological measurements of the plants as reliable methods of direct observation to monitor the “vital signs” of the plants (65,66). Figure 4 (D panel) shows this first stage with the predominance of ecophysiology methods, in which definitions of physiological factors and vegetation damage, pollution prevalence, pests and nutrients appear as drivers of forestry research in the literature (67,68).

On the other hand, the spatio-temporal perspective of forest health is currently under development, constantly incorporating new methodologies mainly focused on the “massive data approaches” at spatial level. These massive data monitoring tools does not only refer to the use of remote sensors for landscape scale assessment, but also other methodologies developed in the last decades in field of ecophysiology and the molecular biochemistry, such as the assessment of gas fluxes at ecosystem level (Eddy co-variance towers), the use of high throughput molecular techniques for microbial communities evaluation or genomic approaches at individual and community levels (e.g., soil proteome, biogechemical cycling, etc…) (69,70). Forest modelling, GIS and remote sensing are needed to manage efficiently and in a sustainable way forest resources (71). These methods allow us to explore the spatial dimension of forest health. In turn, forest modelling allows us to explore the temporal dimension of forest health via long-term and short-term forecasting processes. All these modeling methods have particularly increased in the last two decades becoming especially useful to improve forest management in areas with scarce economic resources.

### 4.4. The way forward: future vision for the forest health concept

In this section we envision how forest health research might evolve in the coming years based on similar disciplines and the gaps found. First, we did not find any article combining the idea of forest health with the concept of “essential biodiversity variable” (EBV) (72). This is one of the most prolific frameworks in the last decade for ecosystem monitoring, but it has not been found among the relevant keywords of our analysis. We believe that the research field of forest health would be very benefited from embracing the EBV framework, especially when considering the description of forest condition. Using EBVs to describe forest health can be useful to increase the comparability of studies carried out in different places.

Similarly, the term “ecosystem services” is also missing in the forest health literature. This concept was introduced in the scientific literature several decades ago (73,74), but it seems to have gone unnoticed in the “forest health” research field (75). We believe that the link between “healthy forests” and their capability to provide ecosystem services might emerge as a new and interesting field of research. While EBVs can help to homogenize the ways of assessing forest health, ecosystem services can contribute to standardizing how we quantify the outcomes provided by forests.

Finally, to integrate the current meaning of the target concept and to transcend it using the above-mentioned proposals, we envision a conceptual and operational alignment between the concepts of “forest health” and “one health” (76). The concept of one health has reached a very holistic meaning in the present time. It used to be focused on single aspects of the health: pain, infection, symptoms, etc. The current meaning put the focus on the concept of health far beyond the absence of illness. One health aims to put together human, animal, and environmental health. This holistic view is slowly moving from the academia into practice (77). This process requires to increase our efforts in transdisciplinary collaboration (78).

Despite the large number of scientific articles related to forest health, the initial literature review found that although they use the term, few authors dare to give a clear and comprehensive approach in their manuscripts. After what we have learned, we recognise that this is a qualitative concept that encompasses the overall state of a complex system studied from various disciplines. According to this holistic approach, we might agree in defining forest health as the capacity of a forest to sustainably provide a wide range of ecosystem services while maintaining biodiversity, natural rhythms and resilience to disturbances inherent in forest dynamics (79–82). Therefore, whether a forest is healthy or not will depend on its natural functioning, buffering capacity and resilience, for which integrated monitoring and management with a vision of conservation of the vital constants and functions of the system is essential. As forest ecosystems are complex systems, assessing and understanding the totality of their functioning is a constant challenge. This means that research on forest health continues to change according to the knowledge needs and concerns observed in forests by scientists and experts, according to available techniques and technologies, policies and social concerns, and the availability of resources, mainly.

## 5. Conclusions

Forest health research has experienced exponential growth in the number of authors and publications, reflecting its relevance and dynamism within the scientific community. We have found a geographical bias in knowledge creation and research focus, as it does not align with the countries that host the largest percentage of the world’s forests (such as tropical countries). This disparity between the Global North and the Global South raises concerns about the integrity and inclusiveness of this field of study.

Concepts and research in forest health demonstrate an evolution towards integration of knowledge over time, and global environmental challenges. Keyword analysis revealed a thematic paradigm shift from the effect of air pollution to current interests in climate change impacts, tree mortality and drought, increasingly integrated with remote sensing technologies and specialised topics such as invasive species and ecophysiology.

The temporal analysis of clustering by descriptors reveals a transition towards complexity and multidisciplinary approaches, showing an evolution from mono- to multi-causal factors. This reflects an effort to understand interactions in complex systems. The integration of advanced methodologies, including remote sensing, forest modelling and big data analysis, has significantly improved the capacity for forest monitoring and management.

Finally, we envisage a conceptual and operational alignment between the concepts of “forest health” and “one health”. The holistic “one health” perspective, which integrates human, animal, and environmental health, can provide a comprehensive approach to forest health. Transdisciplinary efforts and collaboration are needed to bridge the gap between academia and practical implementation.

## Contributions

CAM: Conceptualization, Methodology, Investigation, Formal analysis, Data Curation, Writing - Original Draft, Writing - Review & Editing, Figures 1,2,3,4,6 and Supplementary Material, Supervision; RMNC: Conceptualization, Writing - Review & Editing; FJBG: Conceptualization, Resources, Writing - Original Draft, Writing - Review & Editing; FJRG: Writing & Review; PGM: Conceptualization, Methodology, Investigation, Formal analysis, Writing - Original Draft, Writing - Review & Editing, Figure 5, Supervision.

## Acknowledgments

Thanking Pablo Salazar-Zarzosa for his detailed reading and constructive comments for improvement.

## Supplementary Materials

Table S1: Clustering of the main keywords of the publications search results for forest health, Fig S1: Evolution of the annual scientific production related to Forest Health, Table S2: Main information of the results of the bibliographic search in Web of Science related to Forest Health (Bibliometrix R information), Fig S2: The most relevant 20 authors with the highest number of publications and their evolution of the scientific production evolution, Fig S3: Spatial distribution of the number of publications in different countries on forest health 1934-12/2023 (Graph R from Bibliometrix), Table S3: Scientific production related to forest health 1934-12/2023 by continent, Fig S4: Spatial distribution of the number of publications in different countries on forest health 1934-12/2023 (Graph R from Bibliometrix), Fig S5: Evolution of scientific production related to forest health in the 5 main countries (Graph R from Bibliometrix), Fig S6: Thematic categories of the journals to forest health 1934-12/2023 (WoS graph), Fig S7: Most relevant funding agencies in forest health studies by number of works supported (WoS), Fig S8: Co-occurrence network to 50 most mentioned keywords in the scientific literature (1934-12/2023) (WoSviewer).

## Data availability

The data used to generate the analysis in this article are fully reproducible. They were extracted from the Web of Science database through the query and time period indicated in the methodology.

